## Supplementary Material for "Neural Mechanisms of Creative Problem Solving - From Representational Change to Memory Formation"

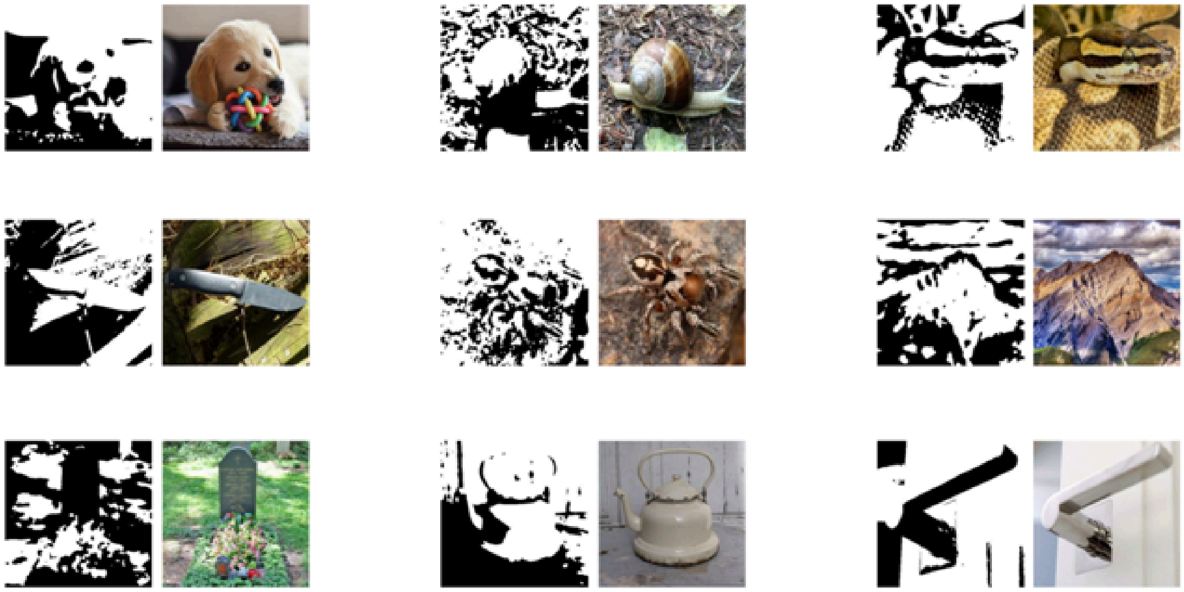

***Figure S1.*** Mooney image examples

*Note*. The black/white abstracted images on the left represent the Mooney images and the corresponding real-world pictures are presented on the right.

### Categories as response options for solution in the MRI scanner

1. Leisure & Hobbies,
2. Tool, weapon or technology,
3. Clothing & Accessories,
4. Vehicles & Road Traffic,
5. Landscape or building,
6. Vehicles & Road Traffic,
7. Predators or rodents,
8. Mammal,
9. Reptiles or Insects,
10. Birds or animal that lives in/near the water,
11. Events or Science,
12. Objects in the house,
13. Human Being

### Relationship between different dimensions of insight experience

We explored whether it is justified to combine the three insight dimensions (*Suddenness, Emotion* and *Certainty*) into one single measure. For this, we calculated a measurement model for a latent insight factor from those three insight dimensions for correctly identified Mooney images. The latent factor was estimated within a confirmatory factor analysis (CFA) in R using the lavaan package (v. 0.6-16, Eves Rosseel, 2012). The model fit was evaluated via Bentler's comparative fit index (CFI), root mean square error of approximation (RMSEA) and standardized root-mean-square residual (SRMR). Accepted thresholds indicating good model fit are RMSEA ≤ .05, SRMR <0.1 and CFI ≥ = .95 (Hu & Bentler, 1998, 1999; Schermelleh-Engel et al., 2014). Note, due to a lack of degrees of freedom no exact *Chi²* goodness-of-fit statistic could be calculated. Error variance of the latent factor was fixed to one to receive factor loadings from all three insight dimensions.

The model converged normally after 12 iterations. The latent insight factor loaded significantly onto all three variables: *Certainty* ($\lambda$=.617, *z*=23.70; *p*<.001), *Emotion* ($\lambda$=.650, *z*=25.17; *p*<.001) and *Suddenness* ($\lambda$=.531, *z*=20.31; *p*<.001) suggesting that those variables contribute significantly to the latent insight factor. Practical fit indices (CFI= 1.0; RSME = 0.000; SRMR =0.000) suggested a good fit of the model to the data.

Note, the significant factor loadings did not change significantly when including all trials (i.e. also incorrectly identified Mooney images). In sum, those results justified combining the three insight dimensions into one single measure for further analyses.

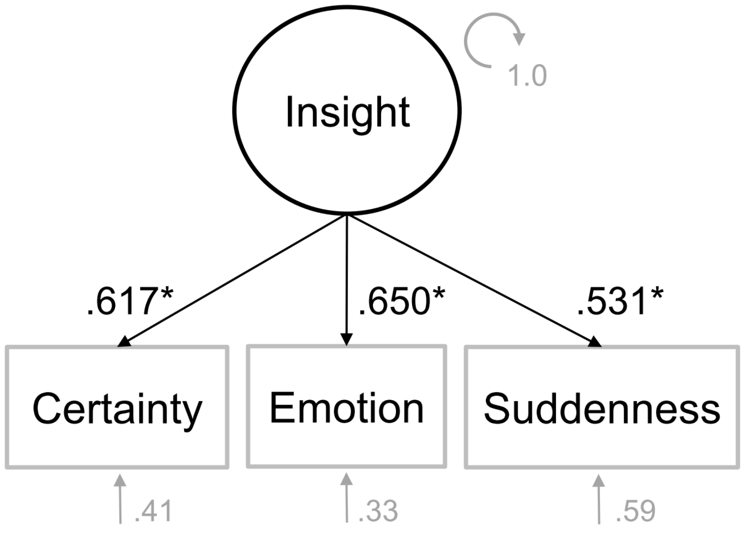


***Figure S2.*** Measurement model for latent insight variable

*Note*. Asterisk = significant loadings at p<.001.

### Estimating threshold for insight trials based on object recognition time

To eliminate trials that might solely involve object recognition and no insight, we excluded all trials with solution times faster than 1.5 seconds (>5*SD of average object recognition time) from our analyses. The time for object recognition was informed by a control experiment, wherein a subset of gray-scaled images (N=78), the same set used to generate the Mooney images, was presented to 40 subjects (19 females, *M*=27.3 years, *SD*= 5.6 years) in an online test recruited via Mechanical Turk and a German student plattform of Humboldt University. Their objective was to press a solution button upon identifying the object, mirroring our Mooney identification task. Median correct object recognition time for these gray-scaled images was 753.2ms (*SD*=140.1ms) (see Fig. S3).

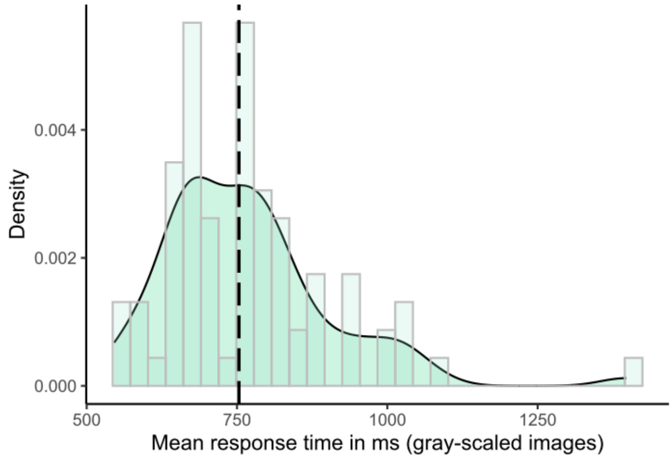


**Figure S3**. Mean response time for object recognition of gray-scaled images.

*Note*. This is a subset (N=78) of those images that were used to create the Mooney images. Line represents median response time.

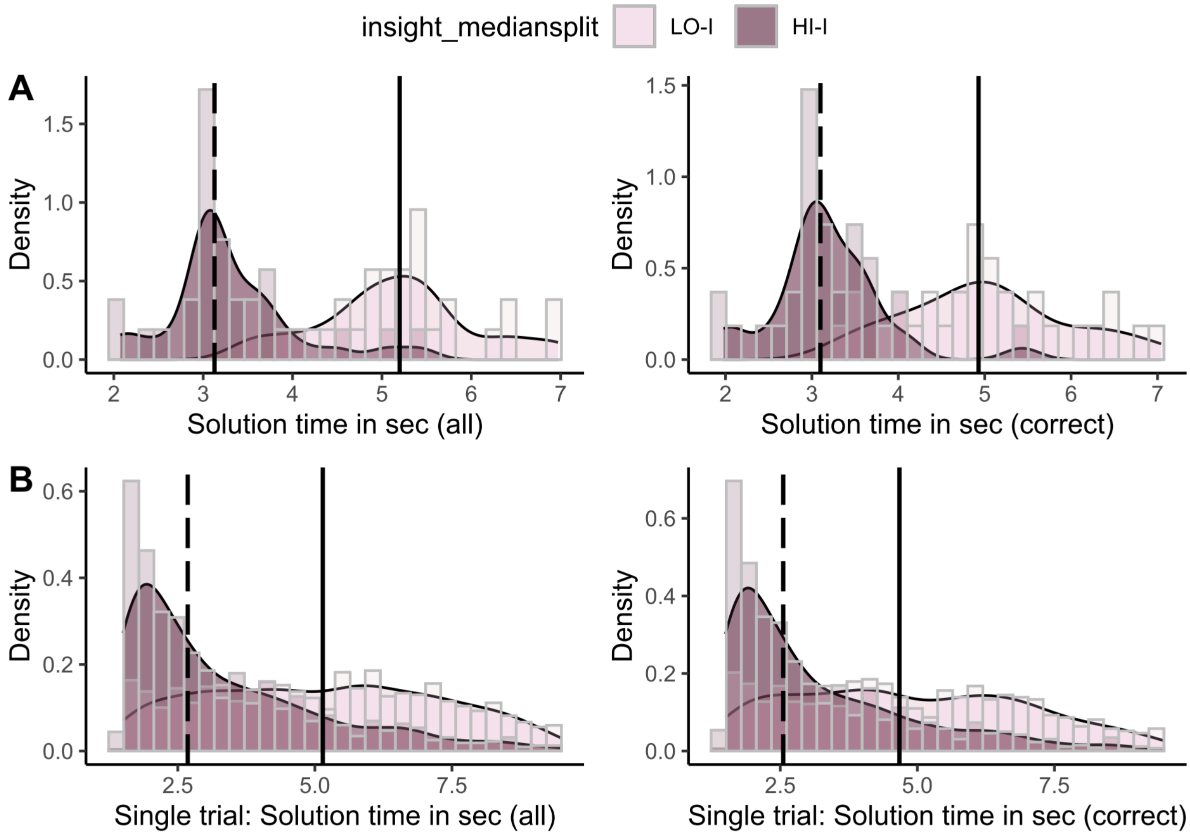


***Figure S4***. Response time distribution divided by high and low insight solutions

*Note*. *LO-I* = low insight trials; *HI-I* = high insight trials. (all) = all solved trials; (correct) = only correctly solved trials. Dashed line = median response time for *HI-I* trials; Full line = median response time for *LO-I* trials. Panel A: Representation of the aggregated solution time distribution across each subject; Panel B: Illustration of the solution time distribution at the individual trial level.

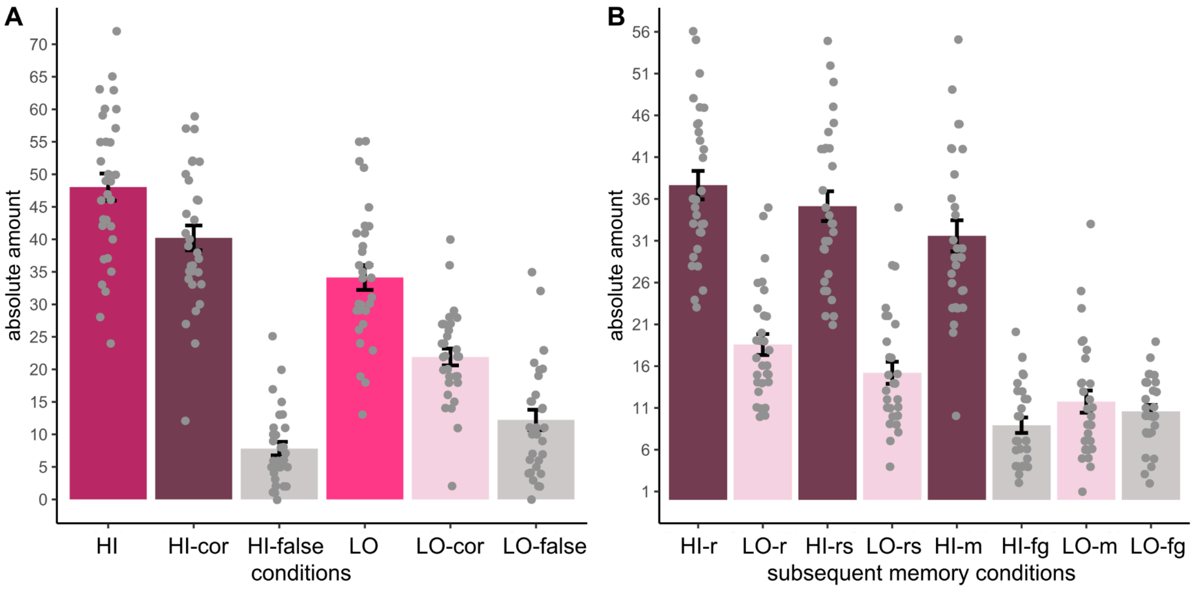

***Figure S5.*** Amount of trials for each insight and insight memory condition

*Note*. **Panel A**: *HI*= high insight trials, HI-cor= correctly solved high insight trials, *HI-false* = incorrectly solved high insight trials, *LO* = low insight trials; *LO-cor* = correctly solved low insight trials; *LO-false*= incorrectly solved low insight trials; **Panel B**: *HI-r* = correctly solved high insight trials & later correctly recognized as having seen before; LO-r = correctly solved low insight trials & later correctly recognized as having seen before; *HI-r*s= correctly solved high insight trials & later correctly recalled as having solved before; LO-rs = correctly solved low insight trials & later correctly recalled as having solved before; *HI-m* = correctly solved and remembered high insight trials, HI-fg = correctly solved & forgotten insight trials, LO-m = correctly solved & later remembered low insight trials, LO-fg= correctly solved and later forgotten low insight trials.

### Univariate fMRI results of VOTC RC areas

For control purposes, aiming to examine whether univariate brain activity in VOTC-RC regions could account for the observed multivariate differences related to insight, we explored the univariate brain activity in pFusG and iLOC in relation to insight. To investigate correctly solved insight trials, we performed the same single trial analyses as for amygdala and hippocampus controlling for run order, solution time and including random subject and item intercepts.

Regarding univariate effects of insight on VOTC-RC areas, there was no significant overall condition effect ($Chi^2(1)=2.96$, $p=.085$, $ß=-.02$) but a condition*ROI interaction ($Chi^2(1)=35.23$, $p<.001$, $ß=.12$) effect. Posthoc analyses revealed a negative effect of insight on BOLD activity in iLOC ($z=-5.51$, $p<.001$, $ß=.07$) but no evidence for an effect in pFusG ($z=1.27$, $p=.20$, $ß=-.40$) (see Fig.S5).

Moreover, we examined potential time-on-task influences on the BOLD signal in both VOTC-RC regions (Yarkoni et al., 2009). To explore this, we replicated the single-trial analyses from above, excluding insight as a predictor. Instead, we computed a solution time*ROI interaction, controlling for run order and incorporating random subject and item intercepts. We found a significant solution time*ROI interaction ($Chi^2(1)=100.38$, $p<.001$, $ß=-.20$). Posthoc analyses revealed a positive effect of solution time on BOLD activity in iLOC ($z=2.87$, $p=.004$) but a negative effect of solution time on BOLD activity in pFusG ($z=-12.11$, $p<.001$).

In summary, the inconsistent univariate effects observed in iLOC and pFusG in relation to insight and solution time make it implausible to attribute the observed consistent multivariate RC effects in both VOTC regions on univariate brain activity.

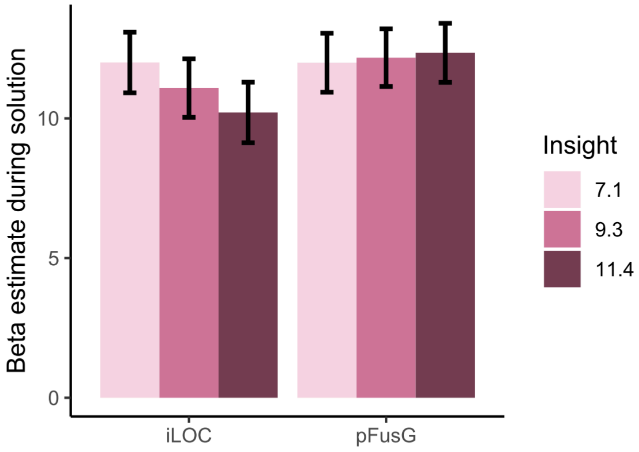


***Figure S6.*** Relationship between univariate BOLD activity in RC VOTC areas and insight.

### Further control analyses to account for the influence of solution time

We conducted additional control analyses to eliminate the potential influence of solution time as the sole factor responsible for the observed effects associated with insight.

*Behavioral analysis*: We adjusted solution time between *HI-I* and *LO-I* trials by excluding fast *HI-I* and slow *LO-I* trials until the time taken to solve both types of trials was no longer significantly different (p>.20). Subsequently, we repeated the mixed model analyses investigating memory as a function of insight (continuous) for correctly solved trials controlling for run (order) and random subject and item intercepts. Odds ratio (OR) and its 95% confidence interval CI) for insight were non-parametrically bootstrapped (1000 resamples) via the *boot* function (v-1.3-30) in R. The results did not change significantly, insight (continuous) still positively predicted memory ($Chi^2(1)=63.83$, $p<.001$, *OR*: 1.576, *CI*[1.41, 1.79]). This result further dismisses differences in solution time as potential explanation for the insight memory advantage.

We also conducted a preregistered control experiment (https://aspredicted.org/xx7hv.pdf) with originally 33 subjects and a final sample of N=27 subjects (70% females, M=23.0 years (*SD*=4.1 years)), recruited through the

Humboldt University student online platform PESA. In this control experiment, participants replicated the same procedures as in the fMRI experiment, using the identical 120 Mooney images. There were two modifications. First, the stimulus disappeared upon pressing the solution button, eliminating potential confounds related to differences in encoding time for the identified Mooney object. Second, the subjects typed their solution instead of selecting a response category which eliminated ambiguity regarding their solution. Participants performed the Mooney identification task in the lab, and five days later (*SD*=0.3 days), they underwent an online subsequent memory test. Six participants were excluded from further analyses due to producing too few correct hits (see preregistration). The data analysis followed the same procedures as for the fMRI sample, with the exception that there was no run (order) variable, as participants completed the experiment in a single run. A distribution of all trials is depicted in Fig. S7-A.

On average, participants found a solution in 45.5% (*SD*=15%) for all presented Mooney images (see Fig. S7-A). The solution was correct in 60.2% (*SD*=17%) of these cases. Of all correctly identified Mooney images, 57% were solved with *HI-I* and 43% with *LO-I* trials (*SD*=12%). Consistent with previous results and the fMRI sample, this *insight-accuracy* effect was significant ($Chi^2(1)=166.0$, $p<.001$, odds ratio=1.55) (Becker, Wiedemann & Kühn, 2020; Danek & Salvi, 2020). The median response time for correct solutions was 4.3sec (*SD*=0.77sec) where participants were faster during *HI-I* (3.6sec, *SD*=0.67sec) than *LO-I* solutions (5.3sec, *SD*=1.0sec). Similar to the fMRI sample, insight significantly predicted response time ($Chi^2(1)=210.43$, $p<.001$, *ß*=-.21). Only accurate responses and response time as covariate of no interest were included into all subsequent memory analyses.

Participants recognized 49.0% (*SD*=16.5%) of the 120 presented Mooney images, regardless of whether they had correctly identified them, with a false alarm rate of 16.3% (*SD*=12.1%). Out of all presented Mooney images, participants correctly solved *and* subsequently correctly recognized 18% (*SD*=9%) and they correctly solved and subsequently correctly recalled having solved them in 15% (*SD*=8%). Finally, out of all presented Mooney images, participants correctly solved and subsequently correctly identified them in 15% (*SD*=8%) (see Fig.S7-B for an overview of a distribution across all conditions). The insight factor (HI-I, LO-I and unsolved trials) significantly predicted variance in subsequent memory

(*Chi²*(2)=113.2, *p*<.001, odds ratio (linear trend for not solved < LO-I < HI-I) = 7.42; see Fig.S7-C). Importantly, when exchanging the continuous insight variable in the mixed model with the binarized insight variable additionally controlling for solution time (excluding unsolved trials), insight still significantly predicted subsequent memory (*Chi²*(1)=25.40, *p*<.001, *odds ratio* = 1.28) (see Fig.S7-D).

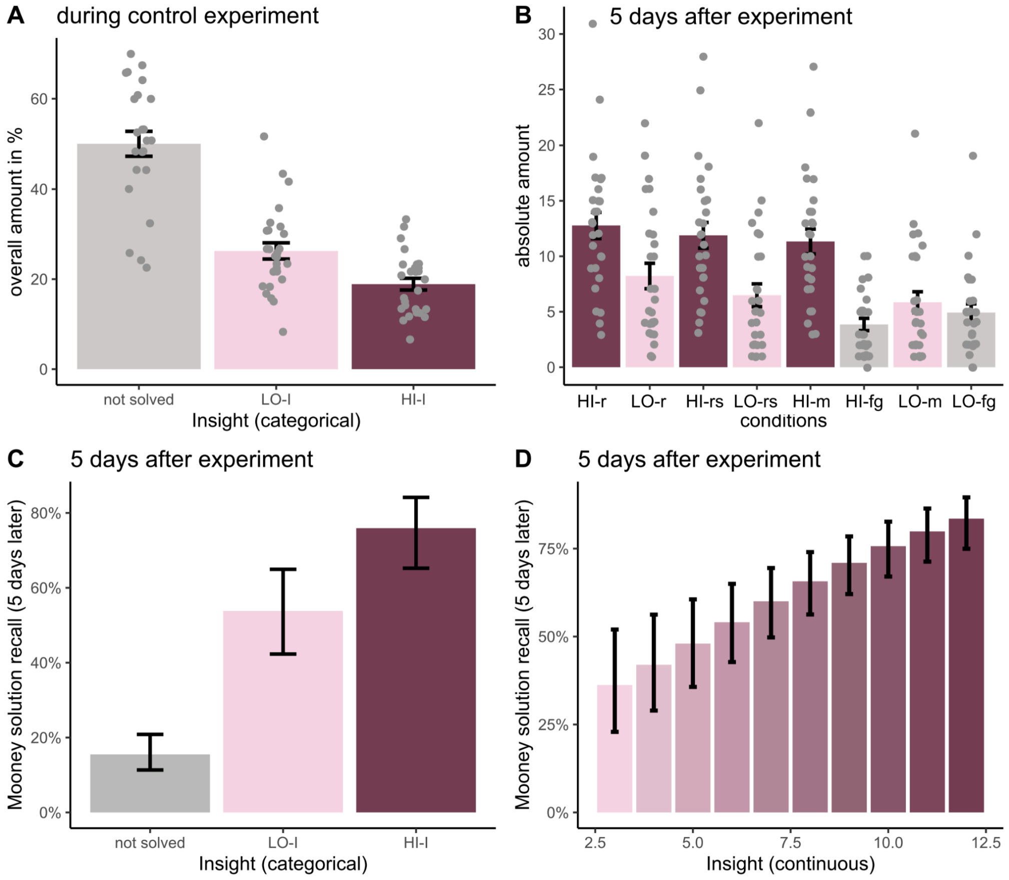


***Figure S7***. Behavioral Insight Memory Effect - control experiment.

*Note*. **Panel A**. overall proportion of all trials per condition; **Panel B**: (remembered = Mooney object correctly identified) *HI-r* = correctly solved high insight trials & later correctly recognized as having seen before; LO-r = correctly solved low insight trials & later correctly recognized as having seen before; *HI-r*s= correctly solved high insight trials & later correctly recalled as having solved before; LO-rs = correctly solved low insight trials & later correctly recalled as having solved before; *HI-m* = correctly solved and remembered high insight trials, HI-fg = correctly solved & forgotten insight trials, LO-m = correctly solved & later remembered low insight trials, LO-fg= correctly solved and later forgotten low insight trials. **Panel C.** Behavioural insight memory effect; *HI-I*= correctly solved high insight trials; *LO-I*= correctly solved low insight trials; estimates are marginal mean percentages to correctly recall the previously identified Mooney image in the respective condition. Error bars represent 95% confidence intervals. **Panel D.** relationship between continuous insight variable (3-12) and memory controlled for solution time; estimates are marginal mean percentages to correctly recall the previously identified Mooney image. Error bars represent 95% confidence intervals.

To summarize, the behavioral control experiment replicated the insight memory advantage observed in the fMRI sample. These findings dismiss longer encoding time of Mooney objects as a potential explanation for the observed insight memory advantage.

*Univariate BOLD analysis*: Similar to the behavioural analysis, we examined the impact of removing fast HI-I and slow LO-I trials on BOLD activity (fast HI-I and slow LO-I trials were removed until the time taken to solve both types of trials was no longer significantly different [p>.20]). Subsequently, we conducted repeated single-trial mixed-model analyses investigating BOLD activity in amygdala and hippocampus as a function of a) insight (continuous) as well as b) insight*memory controlling for run (order) and random subject and item intercepts. To enhance statistical robustness, the p-value was obtained through permutation tests (n = 999) by comparing nested random effects models using the predictmeans package (v.1.1.0) in R (Lee & Braun, 2012). A) insight still significantly predicted BOLD activity during solution ($Chi^2(1)=32.37$, $p=.001$, $ß=.09$) while there was no significant difference between all three ROIs (anterior, posterior hippocampus and amygdala) regarding this relationship ($Chi^2(2)=2.33$, $p=.30$). B) similarly for insight, there was a significant insight*memory interaction ($Chi^2(1)=8.09$, $p=.008$, $ß=.08$) predicting BOLD activity in all three ROIs while there was no evidence for a three-way interaction (ROI*insight*memory: $Chi^2(6)=5.79$, $p=.45$) suggesting no significant difference between the ROIs. A Posthoc analysis revealed that the slope for insight during remembered trials was significantly positive ($z=6.23$, $p<.001$, $ß=.17$) while it was not significantly different from zero during forgotten trials ($z=1.65$, $p=.10$, $ß=.05$).

*Multivariate BOLD analysis*: Making adjustments for solution time in these correlational measures by excluding fast HI-I and slow LO-I trials is not plausible, as each trial represents a correlation of the multivoxel pattern of the current trial in a given ROI with the pattern of every other trial, whether solved fast or slow. However, to account for the potential impact of BOLD activity on the relationship between insight and multivariate measures due to potential time-on-task effects, we conducted a repetition of the aforementioned multivariate RC analyses. This time, we directly incorporated BOLD activity during solution as a covariate of no interest in exchange for solution time additionally controlling for run (order) and random subject and item intercepts. Note, to enhance statistical robustness for the insight-memory analyses, p-values were obtained through permutation tests (n=999) (Lee & Braun, 2012).

Insight remained a significant negative predictor for multivoxel pattern similarity (first RC measure) from pre to post solution (*Chi²(*1)=313.08, *p*<.001, ß=-.14) for both VOTC RC regions when controlling for run (order) and BOLD activity during solution. Posthoc tests showed that pFusG (*Chi²(*1)=211.61, *p*<.001, ß=-.22) as well as iLOC (*Chi²(*1)=155.70, *p*<.001, ß=-.25) reduced their multivoxel pattern similarity from pre to post solution with increasing insight.

Furthermore, the positive insight*time(post solution) interaction predicting representational strength (AlexNet, second RC measure) remained significant (*Chi²(*1)=15.19, *p*<.001, *ß*=.06) for both RC regions (pFusG, iLOC) when controlling for BOLD activity during solution next to run (order) and random subject and item intercepts. There was no evidence that this effect differed between both ROIs (*Chi²(*3)=1.98, *p*=.57). Posthoc analyses revealed that insight negatively predicted representational strength during stimulus onset (*z*=-2.27, *p*=.023) but positively during solution (*z*=2.79, *p*=.005) over both ROIs. Regarding insight-related memory, there was no notable interaction between insight, memory(remembered), and time (post solution) (*Chi²*(1)=0.71, *p-perm*=.40) similar to the outcomes when employing solution time instead of BOLD activity at the solution as a covariate. In contrast, the interaction between memory(remembered) and time (post solution) remained significant (*Chi²*(1)=36.26, *p-perm*=.001, *ß*=.21).

When using Word2Vec as a conceptual model to compute representational strength, the insight*time(post solution) interaction predicting representational strength (second RC measure) remained significant (*Chi²(*1)=16.12, *p*<.001, *ß*=.06) for both RC regions (pFusG, iLOC) when controlling for BOLD activity during solution next to run (order) and random subject and item intercepts. There was no evidence that this effect differed between both ROIs (*Chi²(*3)=1.57, *p*=.66). Posthoc analyses revealed that insight positively predicted representational strength during solution (*z*=3.67, *p*<.001) while there was no evidence for a difference in representational strength as a function of insight during stimulus onset (*z*=-1.55, *p*=.122). Furthermore, there was a significant positive insight*memory(remembered)*time(post solution) interaction (*Chi²*(1)=10.46, *p-perm*=.004, *ß*=.09) predicting representational strength over both brain regions (iLOC, pFusG) while this interaction was not significantly different between both ROIs (*Chi²(7)*=6.63, *p-perm*=.46).

In summary, the outcomes remained largely consistent when incorporating BOLD activity during the solution as a covariate of no interest instead of solution time in the main analyses outlined in the results section.

### Exploratory whole-brain analysis

For exploratory purposes, we also performed a whole-brain analysis to examine whether activity in other brain regions might be parametrically modulated by the intensity of the Aha! experience.

**First-Level Analysis.** We utilized the beta values previously estimated during the first-level analysis (as detailed in the ROI analysis, see Method section). For each participant, simple contrast t-images were generated from the beta weights associated with the onset regressors.

**Second-Level Analysis.** The contrast images representing the parametric modulation of the Aha! experience during the solution phase (button press) were analyzed using SPM's one-sample t-test. Multiple comparisons were corrected by applying a voxel-level threshold of $p<.001$ (Eklund et al., 2016) and a cluster-level threshold of $p < .05$ (family-wise error corrected), and with a height threshold of $t=3.40$. All anatomical regions were identified using the AAL3 atlas (Rolls et al., 2020), based on the percentage of voxels within an activated cluster from the one-sample t-test that intersected the respective anatomical region.

**Whole-Brain Analysis Results:** The intensity of the Aha! experience showed a positive correlation with three distinct clusters (extent threshold k=132 voxels; see Fig. S8). The primary cluster (size: 1,081 voxels, xyz[22, -8, -20]) was located around bilateral hippocampus (covers 32% of this cluster) and extended to bilateral amygdala (11.5%), parahippocampal gyrus (12.5%), ventral striatum (7.5%), olfactory bulb (5.5%), and bilateral putamen (14%). The fact that the largest cluster appeared around the hippocampus and amygdala validates our previous hypothesis that these regions are particularly associated with insight's evaluative component.

The other two clusters were medially situated, with an anterior cluster (size: 341 voxels, xyz[-2, 34, 8]) encompassing the anterior cingulate cortex (70%) and bilateral medial orbital frontal cortex (14%). The ACC has been frequently linked to insight (Becker et al., 2021; for meta-analysis see Dietrich & Kanso, 2010;), and the OFC has also been reported as part of the reward network during insight (Oh et al., 2020). The posterior cluster (size: 458 voxels, xyz[-14, -52, 36]) included the bilateral precuneus (70%), bilateral posterior (15%) and middle (7.5%) cingulate cortex, and the left cuneus (2.5%). The involvement of the precuneus, which is associated with semantic retrieval (Flanagin et al., 2023), is plausible as participants were likely retrieving the semantic content of the object at that moment.


Becker, M., Kühn, S., & Sommer, T. (2021). Verbal insight revisited—Dissociable neurocognitive processes underlying solutions accompanied by an AHA! experience with and without prior restructuring. *Journal of Cognitive Psychology*, *33*(6-7), 659-684.

Dietrich, A., & Kanso, R. (2010). A review of EEG, ERP, and neuroimaging studies of creativity and insight. *Psychological bulletin*, *136*(5), 822.

Eklund, A., Nichols, T. E., & Knutsson, H. (2016). Cluster failure: Why fMRI inferences for spatial extent have inflated false-positive rates. *Proceedings of the national academy of sciences*, *113*(28), 7900-7905.

Flanagin, V. L., Klinkowski, S., Brodt, S., Graetsch, M., Roselli, C., Glasauer, S., & Gais, S. (2023). The precuneus as a central node in declarative memory retrieval. *Cerebral Cortex*, *33*(10), 5981-5990.

Oh, Y., Chesebrough, C., Erickson, B., Zhang, F., & Kounios, J. (2020). An insight-related neural reward signal. *NeuroImage*, *214*, 116757.

Rolls, E. T., Huang, C. C., Lin, C. P., Feng, J., & Joliot, M. (2020). Automated anatomical labelling atlas 3. *Neuroimage*, *206*, 116189.

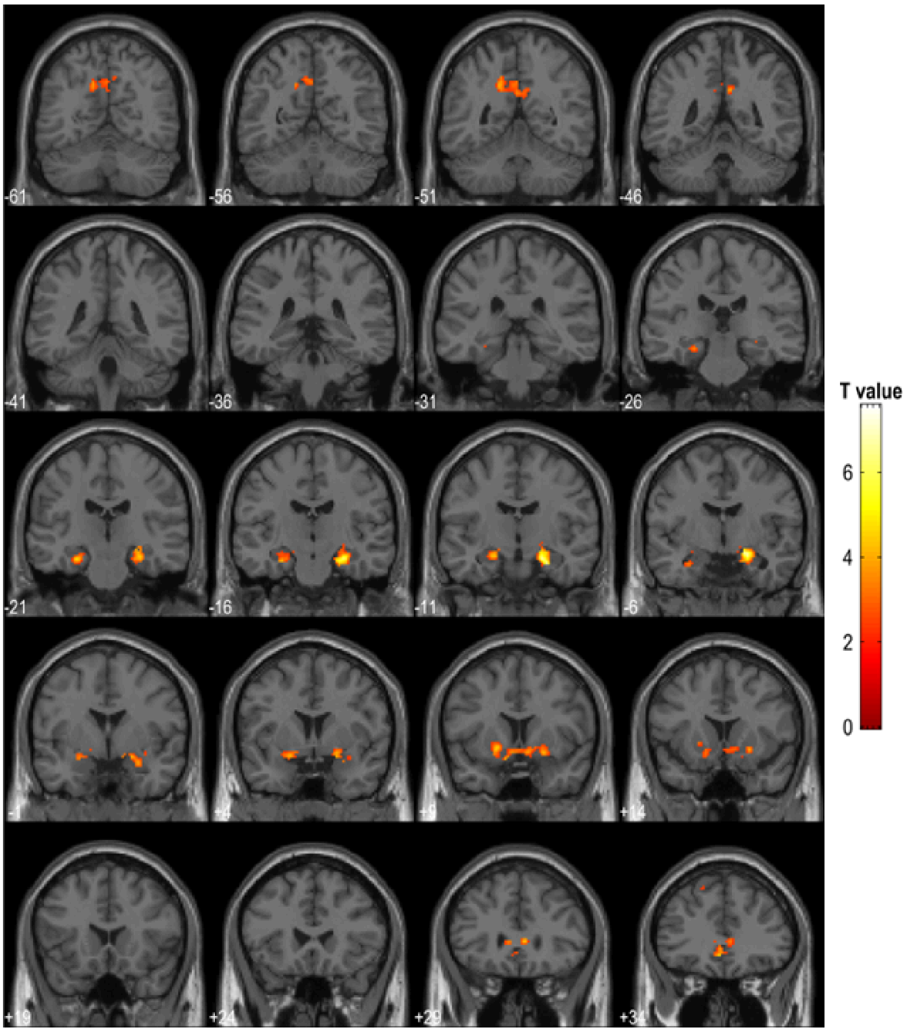


***Figure S8.*** Whole-Brain Activation positively correlated with intensity of Aha! experience

*Note*. Threshold for visualisation: voxel-based threshold p<.001, cluster-threshold p-FWE<.05.

**Table S1.**

Individual Connections from "solution network" for HI-I > LO-I

| ROI1 | ROI2 | T(30) | p-uncor |
|---|---|---|---|
| iLOC l | mITG l | 4.32 | 0.000079 |
| iLOC r | mITG l | 3.46 | 0.000822 |

| | | | |
|---|---|---|---|
| pITG l | mITG l | 3.09 | 0.002136 |
| pFusG r | mITG l | 3.02 | 0.002557 |
| iLOC r | aFusG r | 3.01 | 0.002602 |
| iLOC l | aFusG r | 2.88 | 0.003656 |
| iLOC r | mITG r | 2.76 | 0.004900 |
| iLOC l | mITG r | 2.66 | 0.006275 |
| pFusG l | mITG l | 2.65 | 0.006405 |
| aHC r | aITG l | 2.61 | 0.007060 |
| pFusG l | pITG r | 2.54 | 0.008280 |
| pHC l | aITG l | 2.33 | 0.013467 |
| iLOC l | pFusG l | 2.32 | 0.013530 |
| pFusG l | mITG r | 2.27 | 0.015252 |
| pFusG r | pITG r | 2.25 | 0.015915 |
| aHC r | Amygdala l | 2.23 | 0.016675 |
| pFusG r | aHC l | 2.21 | 0.017485 |
| pFusG l | aITG l | 2.15 | 0.019711 |
| pITG r | mITG l | 2.11 | 0.021769 |
| pFusG r | aITG l | 2.04 | 0.024871 |
| pFusG r | aFusG r | 2.01 | 0.026689 |

| | | | |
|---|---|---|---|
| iLOC r | Amygdala r | 2.01 | 0.026762 |
| mITG r | aITG l | 1.97 | 0.029360 |
| iLOC l | Amygdala l | 1.95 | 0.030032 |
| iLOC l | Amygdala r | 1.91 | 0.032553 |
| aHC l | aITG l | 1.88 | 0.034908 |
| aFusG r | Amygdala r | 1.85 | 0.036759 |
| aFusG l | mITG l | 1.82 | 0.039460 |
| pFusG r | Amygdala r | 1.82 | 0.039621 |
| iLOC r | aHC r | 1.80 | 0.040669 |

Note. The "solution" network contains the ROIs a/pFusG, iLOC, a/m/p ITG, Amygdala, aHC and pHC.

### Graph measures for characterising insight-related more efficient information integration within the *solution network*

*Degree* serves as a reliable indicator of centrality for both networks and ROIs, capturing the level of local interconnectedness within a graph or the entire network. At the ROI level, *Degree* quantifies the edges connected to/from each ROI, while the network degree calculates the mean degree across all graph nodes (Whitfield-Gabrieli & Nieto-Castanon, 2012).

Functional integration within the brain denotes its capacity to rapidly combine specialised information from distributed brain regions, thus illustrating the fluidity of inter-regional communication (Rubinov & Sporns, 2010). Metrics indicating functional integration or the extent of overall interconnectedness in the network comprise *Average path distance* and *Global efficiency*. The *Average path distance* between pairs of ROIs in a graph refers to the minimum connections to navigate between them along an optimal route. Consequently, the node-specific average path distance pertains to the mean path distance between the node and all other nodes in the connected subgraph. *Global efficiency* is calculated as the average of inverse-distances between a given ROI and all other ROIs within the same graph (Whitfield-Gabrieli & Nieto-Castanon, 2012).

Functional segregation within the brain refers to its capacity for specialized processing to unfold within densely interconnected clusters of brain regions (Rubinov & Sporns, 2010). Indicators of functional segregation or localized integration encompass the *Clustering coefficient* and *local efficiency*. The *Clustering coefficient* is delineated as the ratio of interconnected edges in the nearby sub-graph corresponding to each ROI. Node-specific *Local efficiency* is defined as the Global efficiency within the neighboring sub-graph of the given ROI (Whitfield-Gabrieli & Nieto-Castanon, 2012).

**Table S2.**

Mathematical definitions of the used graph measures as implemented in CONN

| Function | Graph measure | ROI-level | Network-level |
|---|---|---|---|
| Centrality | Degree (d) | $d_i = \Sigma_j A_{i,j}$ | $d = \frac{\Sigma_i d_i}{N}$ |
| Functional integration | Average path length (L) | $L_i = \frac{\Sigma_{j\in\Omega_i} D_{i,j}}{N_i - 1}$ | $L = \frac{\Sigma_i L_i}{N}$ |
| | Global Efficiency (GE) | $GE_i = \frac{\Sigma_{i\neq j} 1/D_{i,j}}{N-1}$ | $GE = \frac{\Sigma_i GE_i}{N}$ |
| Functional segregation | Local efficiency (LE) | $LE_i = \frac{\Sigma_{j,k\in\Gamma_i} 1/Dy_{j,k}(i)}{d_i(d_i - 1)}$ | $LE = \frac{\Sigma_i LE_i}{N}$ |
| | Clustering Coefficient (CC) | $CC_i = \frac{\Sigma_{j\neq k\in\Gamma_i} A_{j,k}(i)}{d_i(d_i - 1)}$ | $CC = \frac{\Sigma_i CC_i}{N}$ |

Note. A = adjacency matrix; N = total number of nodes in a graph; di= degree of a graph at each node; D = shortest-path distance matrix, Dy is the shortest-path distance matrix within the neighboring sub-graph at each node, characterized by all nodes neighboring this node and all existing edges among them.

**Relationship between insight-related efficient information integration and subsequent memory**

Finally, for reasons of completeness, we further explored whether different graph measures in the pre-defined *solution network* also predicted insight-related better memory. For this, we repeated FC and graph theoretical analyses but instead of a simple *HI-I>LO-I* contrast, we now calculated an insight (*HI-I vs LO-I)* x memory (remember vs. forgotten) interaction with four conditions: 1) high insight - remembered trials [*HI-I-Rem*], 2) high insight - forgotten trials [*HI-I-Forg*], 3) low insight - remembered trials [*LO-I-Rem*] and 4) low insight - forgotten trials [*LO-I-Forg*] to construct the adjacency matrices. To calculate an interaction, we used the following contrast 1*HI-I-Rem - 1*HI-I-Forg - 1*LO-I-Rem + 1*LO-I-Forg for subsequent FC and graph theoretical analyses. However, please note that due to the low trial count and resulting reduced statistical power, these results should be considered only exploratory.

*Functional connectivity*. The *solution network* did not survive multiple comparisons at the cluster-threshold for an insight x memory interaction.

*Graph theoretical measures*. Global efficiency was significantly increased for this interaction ($t(30)=1.74$, $p<.05$ (one-tailed), $ß=.03$) at the *solution network*-level and also right iLOC ($t(30)=1.70$, *p-uncor*<.05, $ß=.03$) next to left ($t(30)=3.59$, *p-FDR*<.05, $ß=.14$) and right aITG ($t(30)=1.94$, *p-FDR*<.05, $ß=.06$) exhibited significantly more global efficiency compared to the other ROIs in the network. In addition, the insight*memory interaction in the *solution network* was associated with a trend towards significance for increased local efficiency ($t(22)=1.45$, $p=.08$, $ß=.03$). No individual ROI survived multiple comparisons for local efficiency. Neither degree (p>.73), average pathlength (p>.97) nor clustering coefficient (p>.15) showed significant effects in the hypothesized direction.

In sum, the results from FC and graph theoretical measures provide no consistent evidence but some graph measures (increased local efficiency, trend towards significance for local efficiency) suggest an association between insight-related efficient information integration and better subsequent memory.
